# Immunoglobulin A Antibody Composition Is Sculpted to Bind the Self Gut Microbiome

**DOI:** 10.1101/2020.11.30.405332

**Authors:** Chao Yang, Alice Chen-Liaw, Thomas M. Moran, Andrea Cerutti, Jeremiah J. Faith

## Abstract

Despite being the most abundantly secreted immunoglobulin isotype, the reactivity of IgA antibodies towards each individual’s own gut commensal bacteria still remains elusive. By colonizing germ-free mice with defined commensal bacteria, we found the binding specificity of bulk fecal and serum IgA towards resident gut bacteria resolves well at the species level and has modest strain level specificity. IgA hybridomas generated from lamina propria B cells of gnotobiotic mice showed that most IgA clones recognized a single bacterial species, while a small portion displayed polyreactivity. Species-specific IgAs had a range of strain specificities. Given the unique bacterial species and strain composition in each individual’s gut, our findings suggest the IgA repertoire is uniquely shaped to bind our self gut bacteria.

## INTRODUCTION

Immunoglobulin A (IgA) is the most abundant immunoglobulin isotype and plays an essential role ins maintaining the homeostasis of mucosal linings and other physiological processes (*1–4*). In humans, monomeric IgAs mainly exist in systemic circulation, while secretory IgA (SIgA) dominates in secretions, such as stools, saliva and colostrum where it functions as a first line of barrier against pathogens and other environmental insults (*5*). SIgA can aggregate pathogenic bacteria to limit microbial motility leading to reduced invasion into host epithelial cells in the gut (*6, 7*), regulate gut bacteria colonization or clearance through binding bacterial surface epitopes (*8, 9*), and selectively coat disease-associated bacteria in the stool of patients (*10–12*). The production of SIgA is dependent on gut microbiota colonization with much less IgA present in the feces of germ-free (GF) mice than specific pathogen free mice (*13, 14*). Furthermore, monocolonization of GF mice with different bacterial strains leads to increased production of SIgA (*15–18*), although it remains unclear whether the same strains also function as dominant inducers in the context of a complex microbiota.

Considering that SIgA influences gut immune homeostasis in addition to shaping the topography and functions of gut bacteria (*19–21*), it is of paramount importance to better understand the reactivity of SIgA towards gut commensal bacteria, including those that are either colonized on the host (“self” bacteria) and those that are not currently colonized on the host (“non-self” bacteria), such as bacteria from other species or strains colonized in other individuals (*22, 23*). A few studies have used non-overlapping bacteria species to investigate the specificity of gut IgA toward commensals. By employing a reversible germ-free colonization system, Hapfelmeier and colleagues (*24*) observed a long-lived and highly specific SIgA antibody upon colonization of GF mice with *Escherichia coli* (*E. coli*). This SIgA persisted for weeks, even after *E. coli* was decolonized and the mouse returned to the germ-free state. Intriguingly, when new bacteria were then introduced to these pre-colonized mice, the *E. coli*-binding SIgA decreased significantly. Similarly, the gut IgA sequence repertoire of mice colonized with a given microbiota is largely stable but undergoes rapid transformation upon manipulation of the gut microbiota by fecal microbiota transplantation (*25*). This finding implies that the gut IgA repertoire remains malleable and in fact can undergo profound changes in response to modifications of the gut microbiota. Studies looking at the reactivity of gut-derived monoclonal antibodies in mouse or human indicate that these antibodies can recognize specific bacterial strains or species (*26–28*). Some of these gut-derived monoclonal antibodies can also show polyreactivity, which involves recognition of a broad spectrum of bacteria combined with the binding to structurally different but very common microbial or autologous molecules, such as lipopolysaccharide (LPS), double stranded DNA (dsDNA) and flagellin (*29*). These earlier studies have largely focused on simple monocolonized mice or mice colonized by complex microbiotas with well-established bacteria-host interactions. However, the reactivity of native intestinal antibodies or gut-derived recombinant monoclonal antibodies from mice colonized by defined communities of bacteria is still poorly investigated.

To elucidate the specificity of IgA towards the gut microbiota, we dissected the reactivity of both native gut IgA and gut-derived IgA monoclonal antibodies in germ-free mice colonized with diverse, but well-defined bacterial communities. We found that bulk fecal and serum IgA resolved well at the species level with modest strain specificity. By screening the specificity of hybridoma-derived gut IgA clones, we determined that the majority (>75%) of these clones recognized specific bacteria. In addition, these IgA clones retained the capacity of binding bacterial surface antigen in the gut upon oral administration, supporting their potential use as new therapeutics (*7, 26, 27, 30*). The binding-specificities of these monoclonal IgA antibodies largely confirmed our observations for total native IgA with the majority of antibodies resolving at least at the species level and several antibodies being specific towards only the strain of bacteria originally colonized in the gnotobiotic mouse and binding few or no other strains of the same species.

## RESULTS

### Bacteria-induced bulk IgAs have clear species-level resolution and partial strain-level resolution

To explore the specificity of fecal and serum IgA towards the gut microbiota in the context of limited antigenic diversity, we monocolonized C57Bl/6 germ-free mice with one of eight different gut bacterial species with representatives from the most prominent phyla of the human gut including Firmicutes, Bacteroidetes, Actinobacteria and Proteobacteria (Table S1; community 8M) (*31*). We have previously established that monocolonization with each of these strains increases fecal IgA significantly above the low baseline observed in germ-free mice (*17*). After three weeks of colonization to establish steady state IgA levels (*17, 32*), we collected fecal IgA and serum IgA to measure the ability of IgA induced by one bacterial species to bind itself and all seven of the other tested species. The binding capacity of the bulk fecal and serum IgA towards bacterial surface antigens was detected by individually growing each bacterial species *in vitro*, binding each cultured species onto a detection plate, and quantifying antibody binding by ELISA. Interestingly, IgAs from both stool and serum displayed better binding towards species used to colonize mice, relative to those that did not colonize the mice (Fig. 1A), as shown by the prominent signal along the diagonal.

**Fig. 1.**
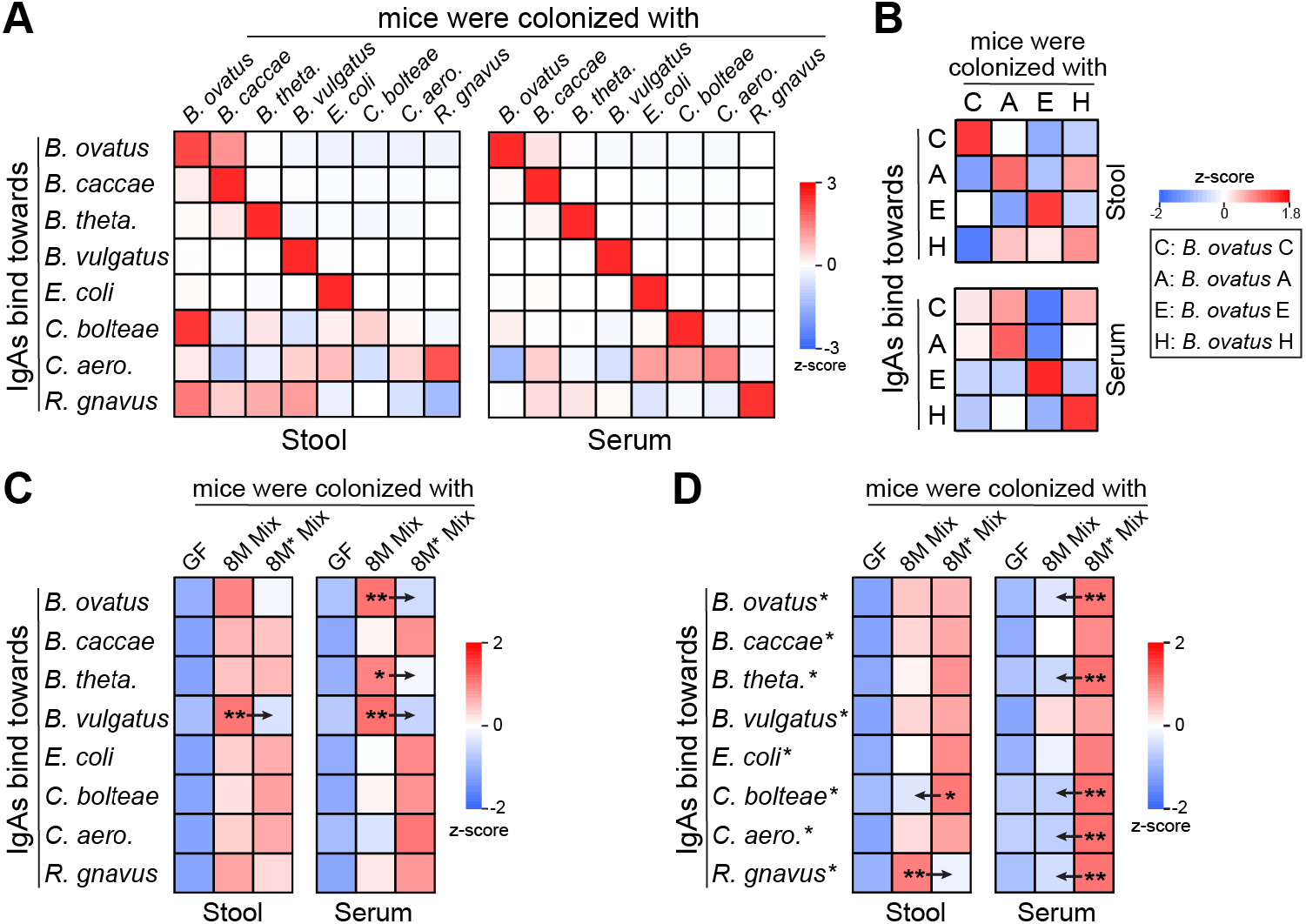
Fecal IgAs bind better towards bacteria that colonized them. (**A**and **B**) Cross-reactivity of stool and serum IgA towards each bacterial species (**A**) or *B. ovatus* strains (**B**). (**C**and **D**) Stool and serum IgA binding capacity towards each bacterial species of a cocktail of 8 bacterial species (8M Mix) or another cocktail of 8 bacterial species, which belong to the same species as 8M Mix but from different strains (8M* Mix). p values were calculated by two-tailed unpaired t test: ^*^p < 0.05, ^**^p < 0.01). Stool and serum samples were harvested from gnotobiotic mice that were colonized with indicated bacterial strains or consortia for three weeks. The average OD_450_ values from 5 to 7 mice were used in heat map plotting. OD_450_: optical density at 450 nm. Detailed information for bacterial strains is listed in table S1. Also see fig. S1.

With the exception of *Bacteroides caccae*, which induced fecal IgA that bound relatively well to *Bacteroides ovatus*, we observed little cross reactivity of bulk IgA even between organisms from the same genus. These results suggest that the majority of the bacteria-induced polyclonal fecal and serum IgA is generated with sufficient specificity to resolve organisms at the species level. Notable exceptions were *C. bolteae* and *C. aerofaciens*, which induced the lowest amount of IgA amongst the eight tested organisms (*17*) and did not bind well to IgA induced from any of the organisms (Fig. S1), and *R. gnavus*, which was highly bound by IgA induced by all eight strains (Fig. S1). This promiscuous IgA binding ability of *R. gnavus* was recently identified to be due in part to the unique ability of this species to bind antibodies that express VH5, VH6, or VH7 variable regions (*33*).

Although unrelated individuals often have substantial overlap in the species composition of their gut microbiomes, there is typically no overlap at the strain level where unique strains of the same species are defined as those differing by at least 4% of their genomic content (*34, 35*). To test if bulk IgA from monocolonized germ-free mice was of sufficient specificity to differentially bind different strains from the same species, we monocolonized germ-free mice with one of four different strains of *B. ovatus* and measured the ability of IgA induced from one strain to bind to each of the other strains of *B. ovatus*. Once again, we found the dominant binding along the diagonal, although with perhaps more off diagonal cross reactivity than was observed on the eight more phylogenetically diverse bacteria tested above (Fig. 1B). These results suggest the specificity of the IgA induced by colonization with a bacterial strain is in part unique to antigens of that particular strain and in part to antigens that are shared across strains from the same species.

To explore the strain level antigen specificity in the context of more complex antigenic diversity, we colonized one set of gnotobiotic mice with all eight bacterial species (8M) from our monocolonization experiments and another set of gnotobiotic mice with a different set of strains (8M*) from the same set of bacterial species (Table S1). Fecal and serum IgA from germ-free controls did not bind well to any of the 16 strains. Although, in general, there was some binding of IgAs from 8M colonized mice to the 8M* strains and vice versa, we observed 11 cases that had significantly different IgA binding between the two strains from the same species. In 10 out of 11 cases, the “self” bulk IgAs better recognized the “self” strain. That is the 8M strain of a given species were bound better by the IgAs from 8M colonized mice, while the 8M* strain of a given species was bound better by IgAs from the 8M* colonized mice. Altogether, these results suggest that the remarkable diversity of the gut microbiota, where every individual has a largely unique set of bacterial strains, drives a similarly unique immune response that is shaped partially by the species composition and partially by the strain composition of each individual. In short, the immune system recognizes “self” microbes better than “non-self”.

### Screening the distribution of lamina propria IgA specificity towards resident bacteria

The above specificities of serum and fecal IgA toward each bacterial strain represent the collective binding of many unique antibodies of different abundance. For more discrete understanding of the species-specific and strain-specific binding capacity of IgA secreted by single IgA^+^ B cells induced by gut bacteria, we made IgA hybridomas using IgA^+^ B cells from the lamina propria (LP) of mice colonized for 3 weeks with the 8M consortia of bacteria. Among all 29 IgA hybridoma clones, the majority of them (93%) expressed a kappa light chain. Like in the bulk IgA reactivity screening, an ELISA was performed to determine the specificity of each antibody towards each of the 8 different strains that were co-colonized in the gnotobiotic mice. Among the 29 hybridoma clones, we found that 8 clones (27.6%) did not bind bacterial surface antigens from any of the 8M community members. Among the 21 bacteria-binding clones, 9 clones (42.9%) bound to only a single strain, 10 clones (47.6%) bound to two species, and 2 clones (9.5%) bound to all the tested bacterial species (Fig. 2A and fig. S2A).

**Fig. 2.**
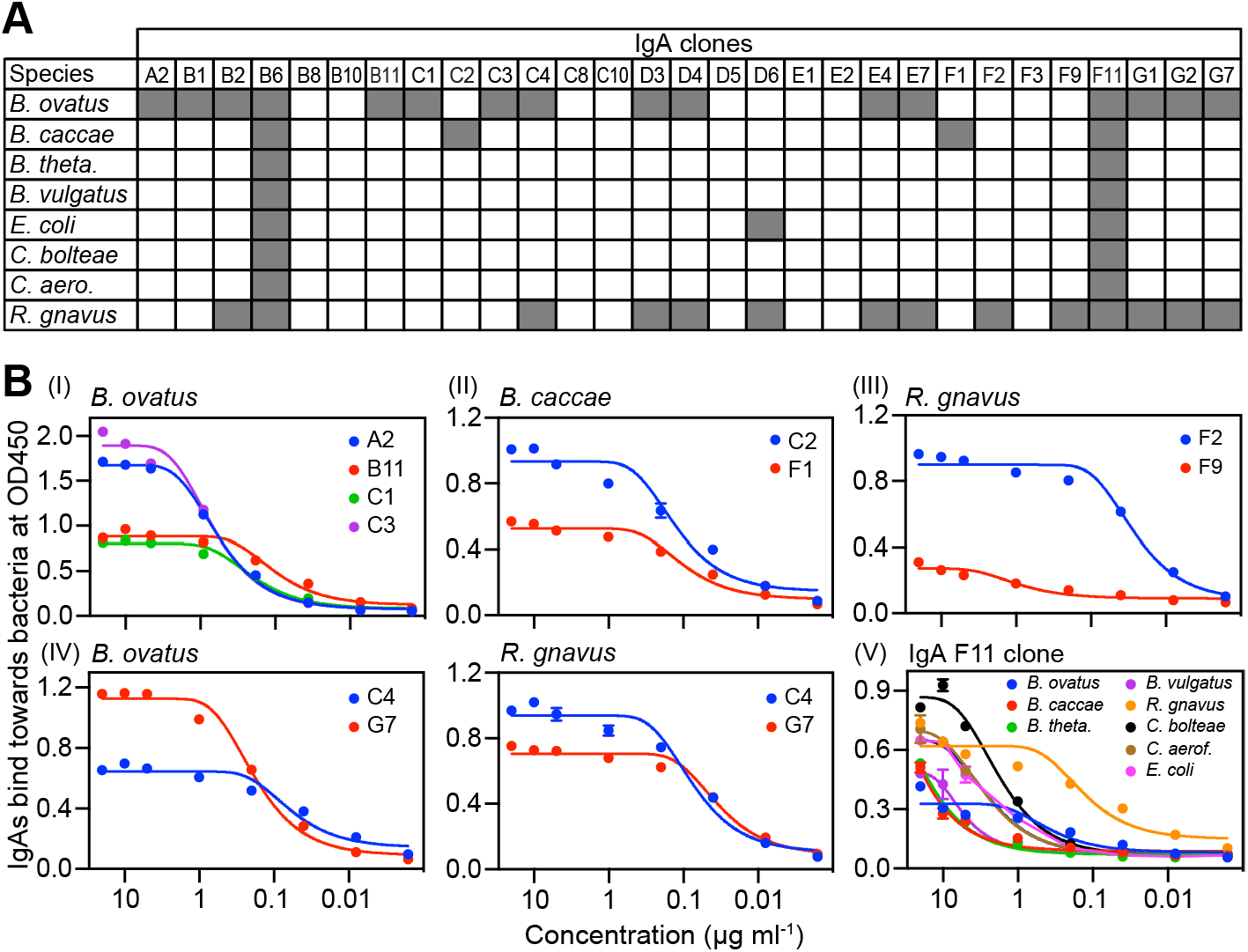
Most monoclonal IgAs isolated from lamina propria IgA^+^ B cells bind bacterial antigens.(**A**) Specificity of 29 monoclonal IgA antibodies towards bacteria that colonized the gut of gnotobiotic mice. Germ-free mice were colonized with all 8 bacterial species for three weeks, then immune cells isolated from intestinal lamina propria were used for hybridoma fusion. Detailed bacterial information is listed in table S1. The OD_450_ > 0.25 at antibody concentration of 10 ug/ml is regarded as positive binding (dark gray square). (**B**) Representative results of hybridoma-produced IgA antibodies against *B. ovatus* (I)*, B. caccae* (II)*, R. gnavus* (III)*, B. ovatus and R. gnavus both* (IV) *and all 8 bacterial species* (V). The relative binding ability of IgA clones was analyzed by ELISA. Monoclonal IgA antibodies after serial dilution were added to ELISA plates, which were coated with each bacterial strain. Results are the average OD_450_ values from two technical replicates. Detailed information of bacterial strains is listed in table S2. OD_450_: optical density at 450 nm. Also see fig. S2.

Similar to findings published previously showing the promiscuous binding ability of *R. gnavus* by IgA antibodies with VH5/6/7 variable regions (*33*), we observed the clones recognizing *R. gnavus* and one other species from the 8M community expressed VH5 or VH6 variable regions (Table S2). Therefore a more accurate interpretation of our results is that 19 out of 21 antibodies (90%) resolved at the species level, which is in line with the species specificity observed in experiments involving total serum or fecal IgA. In addition, we also observed that the IgA clones retained binding capacity towards bacterial surface antigens after passing through the gastrointestinal tract, and could be dosed orally (Fig. S3), which could potentially mimic the naturally produced SIgAs.

The difference in OD_450_ signal of different IgA clones that bind to identical bacteria suggests that these clones recognize distinct epitopes or with different avidity/affinity (Fig. 2B and table S2). Compared to IgA clones recognizing a single bacterial strain from community 8M, the two hybridomas that bound all 8 strains from this bacterial community had a lower overall binding to each microbe as reflected by low OD_450_ value, which may reflect the surface antigen epitopes that are prevalent across diverse taxa (Fig. 2B). We also tested the binding capacity of our IgA clones towards common antigens lipopolysaccharide, double stranded DNA, insulin, flagellin and albumin. Intriguingly, we did not see specific binding to these antigens with mild binding usually observed in promiscuous clones B6 and F11 that could also bound all eight strains (Fig. S2B).

### Screening the specificity of lamina propria IgA towards non-resident bacterial strains

We hypothesized that a fraction of our IgA clones could be strain-specific, or at least unable to bind all strains from a specific species. This could explain our findings of strain-level resolution with polyclonal mixtures from the *in vivo* bulk IgA samples (Fig. 1D). We used five to six additional strains of *B. ovatus*, *B. caccae*, and *R. gnavus* (Table S1) from different human donors to screen the specificity of antibody binding capacity from the hybridomas generated from mice colonized with the 8M community. One of the four *B. ovatus*-binding IgA clones, B11, recognized all six newly tested *B. ovatus* strains (Fig. 3A and fig. S4). IgA clone C1 recognized four out of six strains whereas clone C3 bound two out of six strains. IgA from clone A2 did not bind to any of the new strains. Both *B. caccae-*binding IgA clones only recognize one out of six new strains (Fig. 3B). Finally, the two *R. gnavus* specific monoclonal antibodies bound all five new strains, suggesting they were species-specific (Fig. 3C). We also tested the binding capacity of these bacteria-binding IgA clones towards a larger bacterial library, which included 18 different bacterial species belonging to different phyla. Consistent with our observations, none of the antibodies showed binding to these new species (Fig. S4D-F) confirming their species resolution. Overall these *in vitro* results with monoclonal antibodies demonstrate the majority of monoclonal IgA antibodies resolve at least at the species level. Amongst these species-specific antibodies, the specificity appears to vary from being capable of binding all members of the species to only a subset of strains in the species – presumably driven by the variation of the IgA-binding antigen/epitopes in the species pan-genome. These observations for monoclonal antibodies parallel our *ex vivo* results with bulk serum and fecal IgA antibodies, where the resolution of polyclonal IgA was prominently species-specific and, in some cases, strain-specific.

**Fig. 3.**
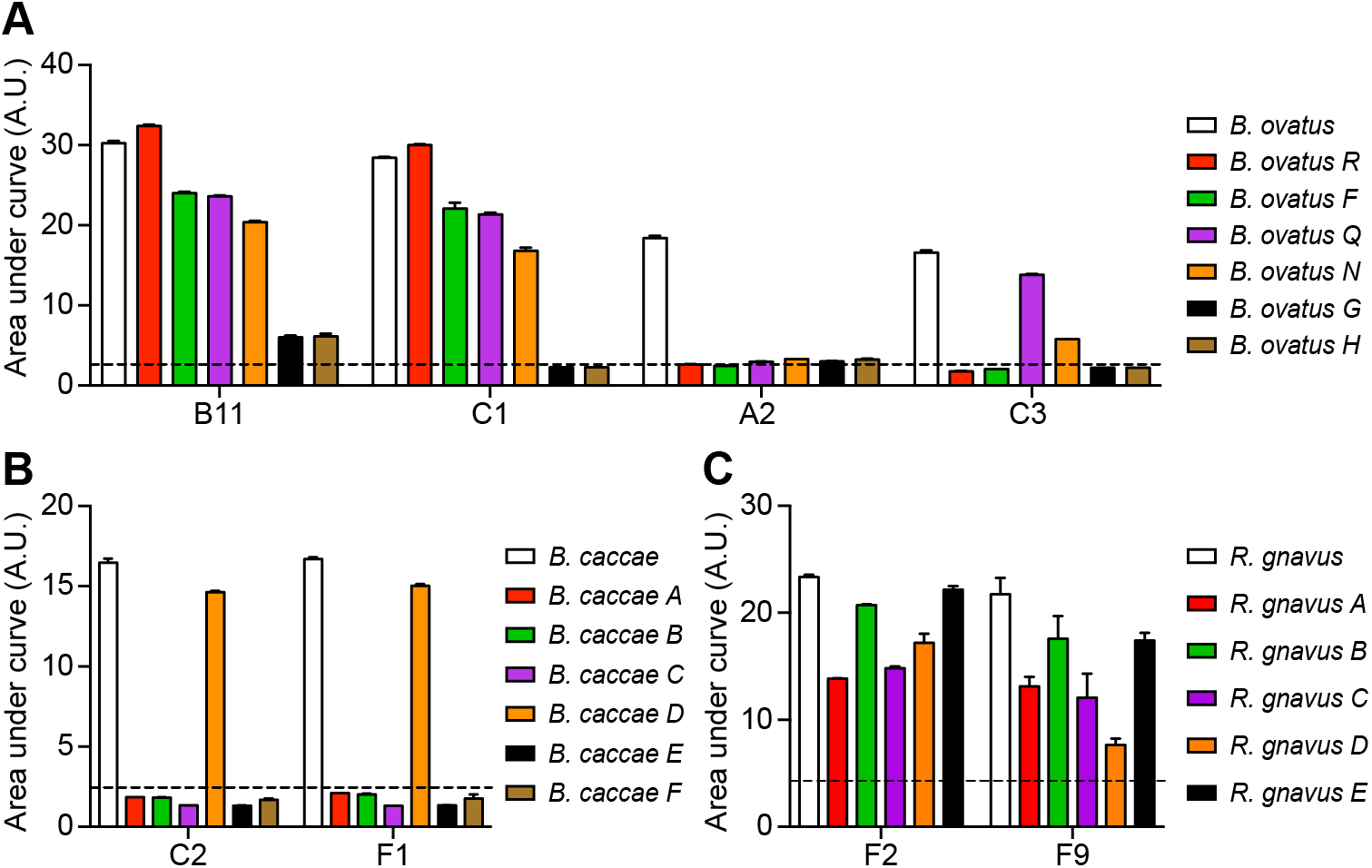
Monoclonal IgAs isolated from lamina propria bind bacteria at species and strain levels. (**A-C**) Specificity of monoclonal IgA antibodies towards different *B. ovatus* strains (**A**), *B. caccae* strains (**B**) and *R. gnavus* strains (**C**) that were were not used to colonize mice. The binding ability of IgA clones was quantified by ELISA. Monoclonal IgA antibodies after serial dilution were added to ELISA plates, which were coated with the indicated bacterial strains. Results are the average OD_450_ values from two technical replicates. Strains denoted by the white/open bar belong to the 8M community that colonized gnotobiotic mice. Detailed information of bacterial strains is listed in table S1. OD_450_: optical density at 450 nm. A.U.: arbitrary units. Dotted lines denote A.U. in negative control group. Also see fig. S3.

## DISCUSSION

Gut IgAs facilitate host-microbe interactions, constrain commensals within the intestinal lumen, modulate bacterial behaviors and shape the composition and topography of bacteria (*9, 11, 36–39*). However, the fine specificity of gut IgAs towards commensal bacteria still remains elusive. Here we demonstrated that bulk IgAs from both stool and serum recognizes “self” bacteria that colonized the host more effectively than “non-self” bacteria of the same species but from a different host. We also observed that about 75% of hybridoma-derived IgA clones generated from single gut LP B cells of gnotobiotic mice resolve bacteria at least at the species level, and only a few IgA clones display polyreactivity. Interestingly, some IgAs bind specifically to the host colonized strain, while others bind to a variable number of strains that belong to the same species but are extraneous to the host.

Several studies have investigated the reactivity of gut IgA towards bacterial surface antigens. Upon monocolonzation, bacteria-reactive IgAs can be detected *in vitro* by ELISA upon culturing the fragments of Peyer’s patches and by ELISpot on culturing purified LP immune cells (*40*). These observations have been later confirmed by cloning IgA-secreting B cells from the LP of monocolonized mice (*32*). In agreement with our results, most IgAs released by single B cell clones from the human gut LP have been shown to react against single bacterial species (*41*). However, it has also been reported that most IgA clones from the LP of specific pathogen-free mice display polyreactivity for common antigens such as DNA, LPS, and flagellin in addition to recognizing a broad range of gut microbes (*29*).

Many factors may account for the discrepancy. First, there may be strain-level differences among bacteria, which implies that specificity screening should be run with carefully selected bacterial strains to distinguish between resident “self” and non-resident “non-self” strains. Second, the degree of complexity of bacterial consortia used to colonize gnotobiotic mice likely has an impact on the production of IgA antibodies. Third, limitations in the duration of colonization in gnotobiotic mice may lead to a relatively increased production of “innate-like” IgA antibodies compared to the largely germinal center-derived IgA antibodies normally found in the gut of an adult mouse with an non-manipulated microbiota. Accordingly, SPF mice are exposed to bacteria much longer than gnotobiotic mice and this prolonged exposure time may underpin the production of “natural” IgA antibodies.

Of note, our results also demonstrate the hybridoma-derived IgAs can transit through the harsh environment of the gastrointestinal tract and retain binding capacity against bacterial antigens. This observation clearly implies the potential usage of *in vitro* engineered IgA antibodies as therapeutics (*7, 42–44*). Compared with naturally secreted IgA antibodies, those hybridoma-derived IgAs lack the bound secretory component, which may finally compromise their stability to bacterial proteases in the gut lumen (*45, 46*). Thus, it would be important to dissect the capability of gut bacteria, including both commensal and pathogens, to degrade secretory component-free IgAs prior to advancing hybridoma-derived IgAs as therapeutics.

In summary, our data show that gut bacteria-induced IgAs have higher reactivity towards “self” strains in mice colonized by a single microbe or a consortium of bacteria. We also demonstrate that the host generates species-specific and strain-specific rather than polyreactive gut IgA antibodies and species-specific IgA antibodies may or may not cross-react with a few to many different strains from the same species. By unraveling the complex specificity of bacteria-induced gut IgA antibodies in gnotobiotic models, our findings highlight the potential role of gut IgAs in the symbiotic host-microbe interactions, because the coating of bacteria by IgA antibodies influences not only the homeostasis of the mucosal immune system but also the physiology of bacteria (*19, 22, 23, 47*).

## MATERIALS AND METHODS

### Mice

Germ-free mice were bred and maintained in flexible plastic gnotobiotic isolators (Class Biologically Clean, Ltd.) in Icahn School of Medicine at Mount Sinai. All mice were housed in groups under controlled temperature, a 12-hour light/dark cycle, and allowed *ad libitum* access to diet and water. Usage of animals for experiments was approved by the Institutional Committee on Use and Care of Animals (IACUC) in Icahn School of Medicine at Mount Sinai.

### Inoculation of germ-free mice with cultured bacteria

In experiments involving gnotobiotic mice usage and colonization, mice were handled as previously described (*17, 35*). Briefly, germ-free mice (~8 weeks old) were transferred outside from isolator into a laminar flow biosafety hood that was pre-cleaned with CLIDOX-S^®^. Frozen bacteria cultures in anaerobic glass vials were sterilized with CLIDOX-S^®^. After being thawed, ~200 µl aliquot of bacteria suspension was introduced into recipient mice via oral gavage. Colonized mice were then housed in clean filter top cages, which contained sterilized mouse diet, water and bedding, for the duration of the experiment.

### Growth of bacterial strains

*B. ovatus* (ATCC 8483)*, B. caccae* (ATCC 43185)*, B. thetaiotaomicron* (VPI 5482)*, B. vulgatus* (ATCC 8482)*, R. gnavus* (ATCC 29149)*, C. bolteae* (ATCC BAA-613)*, C. aerofaciens* (ATCC 25986) and *E. coli* (K-12 MG1655), *P. johnsonii* (DSMZ 18315) and *B. intestinalis* (DSMZ 17393) were obtained from global repositories: the American Type Culture Collection (ATCC) and Deutsche Sammlung von Mikroorganismen und Zellkulturen (DSMZ). Apart from *E. coli*, the bacterial strains were grown under anaerobic condition at 37°C in Brain Heart Infusion (BHI) medium supplemented with 0.5% yeast extract (Difco Laboratories), 0.4% monosaccharide mixture, 0.3% disaccharide mixture, L-cysteine (0.5 mg/ml; Sigma-Aldrich), malic acid (1 mg/ml; Sigma-Aldrich) and 5 µg/ml hemin. LB Broth Miller (EMD Chemicals, Inc.) was used for *E. coli* culture under aerobic conditions at 37°C. After bacterial cultures reached an OD_600_ of 1-2, glycerol was added to a final concentration of 15% (v/v). The mixture was aliquoted into glass vials with a crimped top and stored at −80°C freezer for further use.

### Isolation of single bacterial strains from human stool samples

Except for strains obtained from public culture repositories (ATCC and DSMZ), individual bacterial strains in tables S1 and S2 were isolated from previously banked, deidentified stool samples (*17, 48*). These bacteria were cultured under anaerobic conditions, and the identity of each individual bacterium was verified by mass spectrometry (Bruker Corp.).

### Lymphocyte isolation from mice intestines

Laminar propria lymphocytes were isolated as described previously (*48*). Briefly, small intestines and colons were excised from gnotobiotic mice, followed by cleaning visceral fat and intestinal content. Tissues were opened longitudinally, washed twice in Hank's Balanced Salt Solution (HBSS) without Ca^2+^ and Mg^2+^ (GIBCO) and incubated in dissociation buffer (HBSS including 10% fetal bovine serum (FBS), 5 mM EDTA and 15 mM HEPES) for 30 min at 37°C with agitation to remove epithelium and intraepithelial lymphocytes. The remaining tissues were then washed three times in ice cold HBSS, cut into ~2 cm pieces and digested with collagenase (Sigma-Aldrich), DNase I (Sigma-Aldrich) and dispase I (Corning) for 40 min at 37°C with mild agitation. Cell suspensions were filtered through 70 µm cell strainers, washed three times and re-suspended in IMDM/2%FBS.

### Generation of hybridoma clones

Sp2/0-Ag14 myeloma cell line (ATCC CRL-158) was cultured in Dulbecco's Modified Eagle's Medium (DMEM) supplemented with 10% FBS. Before fusion, 80-90% confluence was achieved. After counting cell number, Sp2/0-Ag14 myeloma cell line and LP lymphocytes were mixed thoroughly and carefully at a ratio of 1:5. Mixed cells were washed in Hybridoma-SFM. Cell pellet was loosened gently. Then polyethylene glycol (PEG) was slowly added along the side of the tube. Tube was incubated at 37°C for 1 minute and then a 10 times volume of Hybridoma-SFM was slowly added to PEG treated cells. After spinning down, the fused cells were incubated overnight in complete media. The next day, the cells were centrifuged and resuspended in semi-solid hybridoma selection medium containing hypoxanthine, aminopterin and thymidine (HAT), mixed well and gently transferred to petri dishes. After incubation at 37°C for 2 weeks, single cell colonies were picked into 96 well plates by colony picker (Hamilton Company) and cultured for another 2 days for antibody isotyping. An aliquot of cell suspension was taken out and frozen for further examination.

### Production of IgA antibody

Selected clones of IgA hybridoma cells were thawn and expanded in DMEM buffer supplemented with 10% FBS (Hyclone). When hybridoma cells grew to 90% confluence, they were washed three times in PBS and then transferred to the lower level chamber of CELLine Disposable Bioreactor (Corning). Hybridoma-SFM (Gibco by Life Technologies) was added to the upper level of the Bioreactor. In the first two weeks, Hybridoma-SFM was changed within 24 hours when medium changed color from red to orange. When a very high density of cells was observed, about 50% of cells and medium in the lower chamber were harvested. After spinning down, supernatant was kept for antibody purification. Cells in the bioreactor were cultured for four weeks and medium in the second level of the chamber were frequently harvested. IgA antibody in medium was precipitated in ammonium sulfate solution and then dialyzed in a cassette in PBS overnight at 4°C. Concentration of dialyzed IgA antibody was quantified by ELISA. Production of larger IgA quantifies for *in vivo* oral dosing was performed by ProSci Incorporated. Briefly, the hybridoma cell line was cultured to 85% confluence and injected to peritoneal cavity of BALB/c mice. Ascites were harvested, precipitated and dialyzed.

### Detection of bacteria-binding IgA antibodies by ELISA

Bacteria were cultured to OD_600_ = 1~2 in appropriate condition and then fixed in 0.5% formaldehyde in PBS for 20 min at room temperature. After washing 3 times, bacteria density was adjusted to OD_600_ = 1 in PBS. ELISA plates were coated with 30 µl of the adjusted bacterial suspension and incubated at 4°C overnight. After washing and blocking with 1% BSA, diluted fecal or seral samples containing polyclonal IgA were added and incubated overnight at 4°C. Captured IgA was detected by horseradish peroxidase (HRP)-conjugated goat anti-mouse IgA antibody (Sigma-Aldrich; A4789). ELISA plates were developed by TMB microwell peroxidase substrate (KPL, Inc.; 50-76-03) and quenched by 1 M H_2_SO_4_. The colorimetric reaction was measured at OD = 450 nm by a Synergy HTX Multi-Mode Microplate Reader (BioTek Instruments, Inc.).

### Screening the specificity of hybridoma-produced IgA by ELISA

For the screening of monoclonal IgA antibodies against bacterial antigens, reciprocal dilutions of hybridoma-produced IgA antibodies were added to ELISA plates that were coated with different whole bacteria as described above. The polyreactivity of hybridoma-obtained IgA was tested by adding them to ELISA plates that were coated with common antigens dissolved in PBS before coating ELISA plates: Lipopolysaccharide (Sigma-Aldrich; L4391), double stranded DNA (Sigma-Aldrich; D4522), insulin (Sigma-Aldrich; 91077C), flagellin (Sigma-Aldrich; SRP8029) and albumin (Sigma-Aldrich; A1653).

### VDJ sequencing of IgA hybridoma clones

Hybridoma cell lines were cultured in DMEM buffer supplemented with 10% FBS. Cells were harvested at 90% confluence after washing three times with PBS. Total mRNA was extracted with the RNeasy Mini Kit (QIAGEN, 74104) according to manufacture’s protocol. cDNA was synthesized with High-Capacity cDNA Reverse Transcription Kits (Applied Biosystems, 4368813) and approximately 200 ng cDNA was used for PCR amplification with Phusion High-Fidelity PCR Master Mix with HF Buffer (ThermoFisher Scientific, F531L). The primers for amplicons of murine IgA are VH promiscuous 5’-GAGGTGCAGCTGCAGGAGTCTGG-3’ in combination with Cα (outer) 5’-GAGCTCGTGGGAGTGTCAGTG-3’ as described previously (*25*). PCR conditions for IgA amplicons were set as follows: 30 s at 98°C, and then 28 cycles of (98°C for 10 s; 62°C for 30 s and 72°C for 45 s); and then 72°C for 5 min. Amplicons were sent for Sanger sequencing to Psomagen, Inc. Sequenced reads were analyzed with ImMunoGeneTics(IMGT) V-QUEST (http://www.imgt.org/) (*49*) using reads from both primers. Only results consistent for both primers were displayed.

### Statistical analysis

Data are shown as mean ± SEM. Statistical significance between two groups was assessed by an unpaired, two-tailed Student’s *t* test. Comparisons among three or more groups were performed using One-way ANOVA. Data plotting and statistical analysis were performed using GraphPad Prism 7.0 (GraphPad Software, La Jolla, CA). A *p*-value less than 0.05 is considered statistically significant.

## ACKNOWLEDGEMENTS

We are grateful to Drs. C. Cunningham-Rundles, E. Grasset, and B. Brown for helpful discussions and comments. Dr. I. Mogno, Z. Li and M. Spindler for help with bacterial isolates. This work was supported in part by the staff and resources of Gnotobiotic Mouse Core Facility and Microbiome Translational Center in Icahn School of Medicine at Mount Sinai. This work was supported in part by NIH National Institute of Diabetes and Digestive and Kidney Diseases Grants DK124133, DK112978, DK124165, and DK123749 and a Crohn’s Colitis Foundation Senior Research Award.

## AUTHOR CONTRIBUTIONS

C.Y. and J.J.F conceived the study and designed the experiments; C.Y., A.CL. and T.M.M. conducted the experiments; C.Y., A.C. and J.J.F. analyzed and interpreted data; C.Y. and J.J.F. wrote the manuscript.

