## Supplementary Information for "Immunoglobulin A Antibody Composition Is Sculpted to Bind the Self Gut Microbiome"

### SUPPLEMENTARY MATERIALS

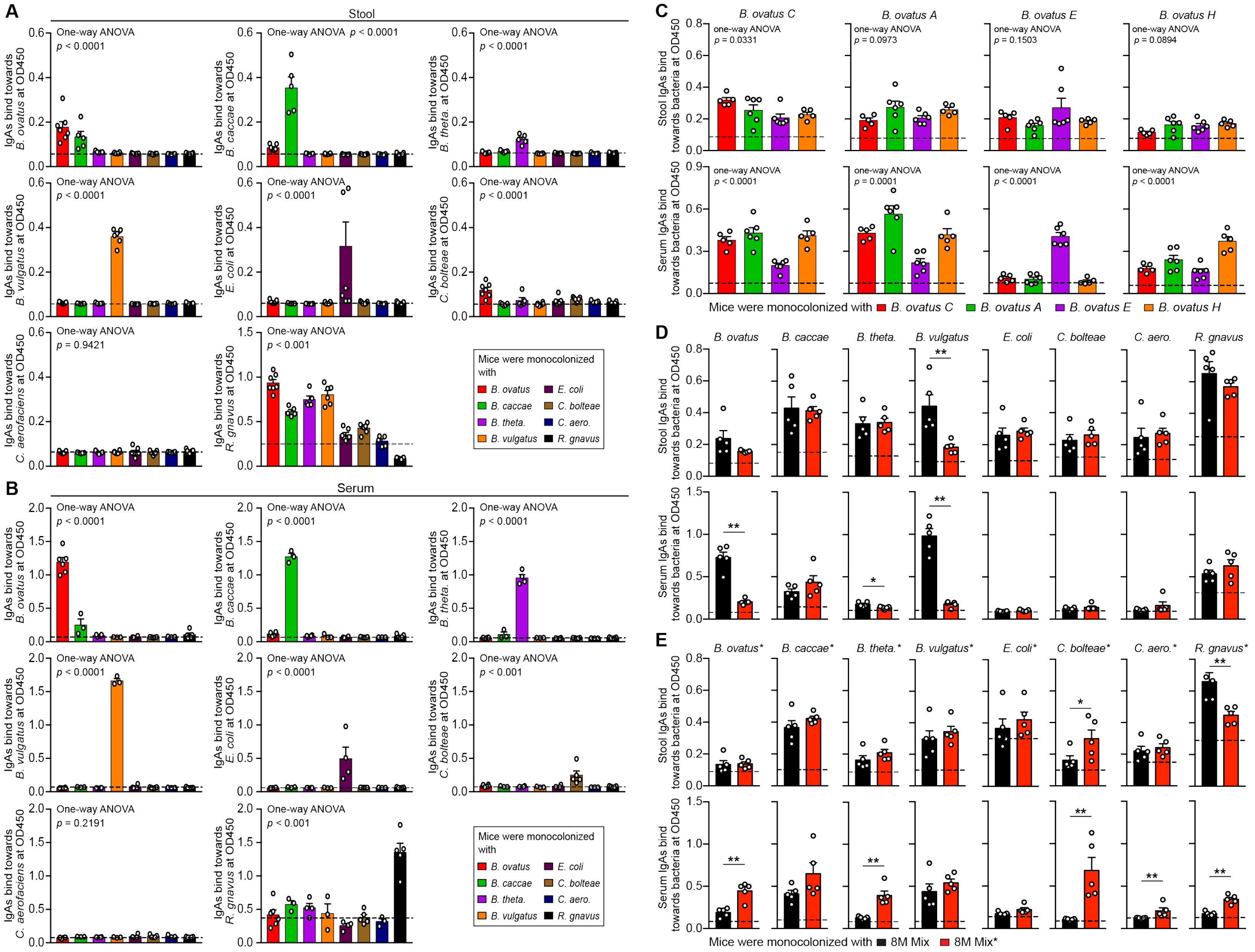

**Fig. S1. Values of fecal IgAs binding towards bacteria at $OD_{450}$, related to fig. 1.** (**A**-**C**) $OD_{450}$ values of stool and serum IgA towards each bacterial species (**A** and **B**) or *B. ovatus* strains (**C**). (**D** and **E**) $OD_{450}$ values of stool and serum IgA towards individual bacterial species of a cocktail of 8 bacterial species (8M Mix) (**D**) or another cocktail of 8 bacterial species (8M* Mix) that belong to the same species with 8M Mix but different strains (**E**). *p* values in A, B, C were calculated with One-way ANOVA; *p* values in D and E were calculated with two-tailed unpaired t test ($^{*}p < 0.05$, $^{**}p < 0.01$). $OD_{450}$: optical density at 450 nm.

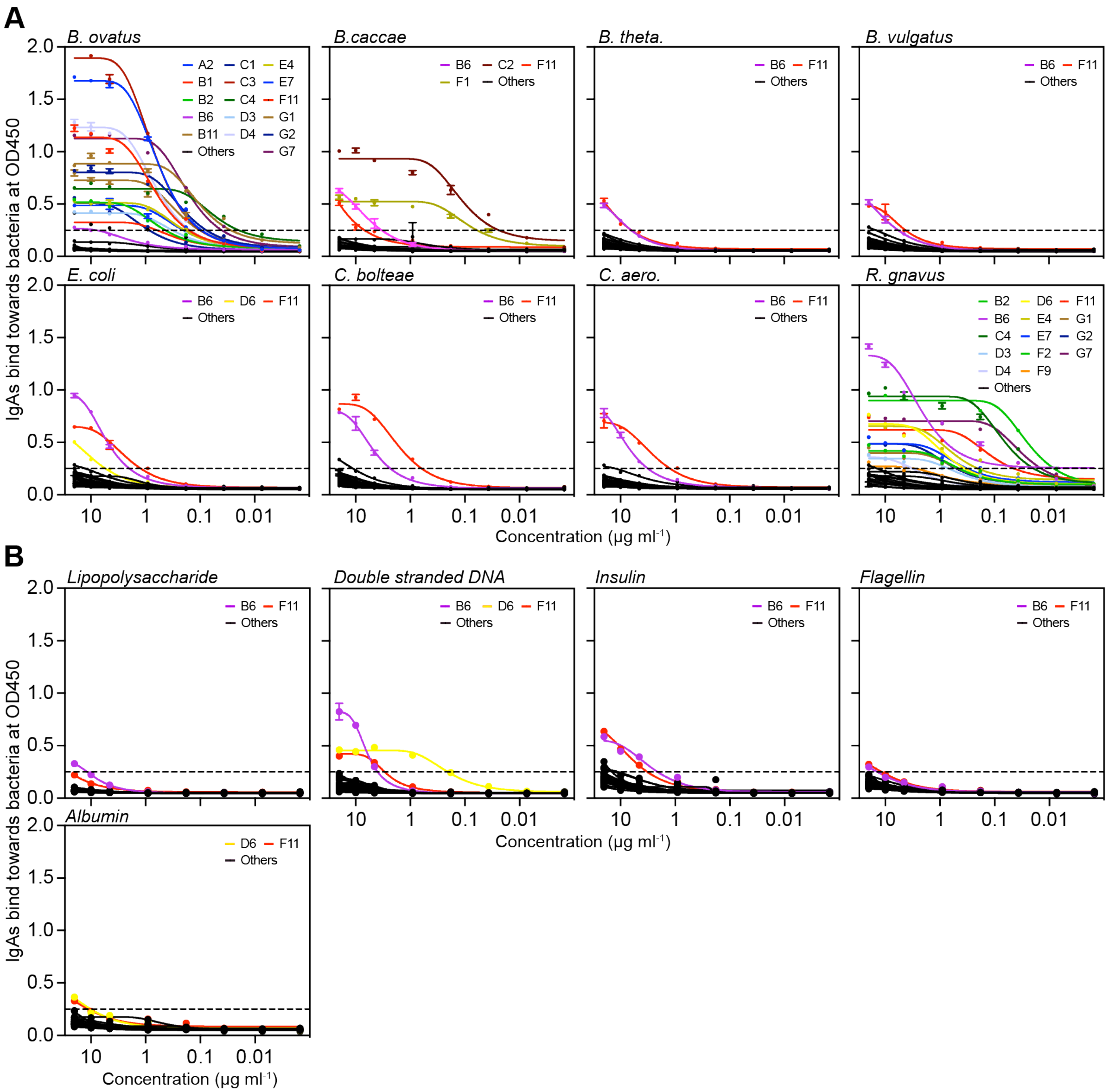


**Fig. S2. Specificity of hybridoma-produced IgA towards bacterial surface antigens and common antigens, related to fig. 2.** (**A** and **B**) ELISA analysis of hybridoma-produced IgA against bacteria that colonized the gut of gnotobiotic mice (**A**) and common antigens including lipopolysaccharides, double stranded DNA, insulin, flagellin and albumin (**B**). Results are the average $OD_{450}$ values from two technical replicates. $OD_{450}$: optical density at 450 nm.

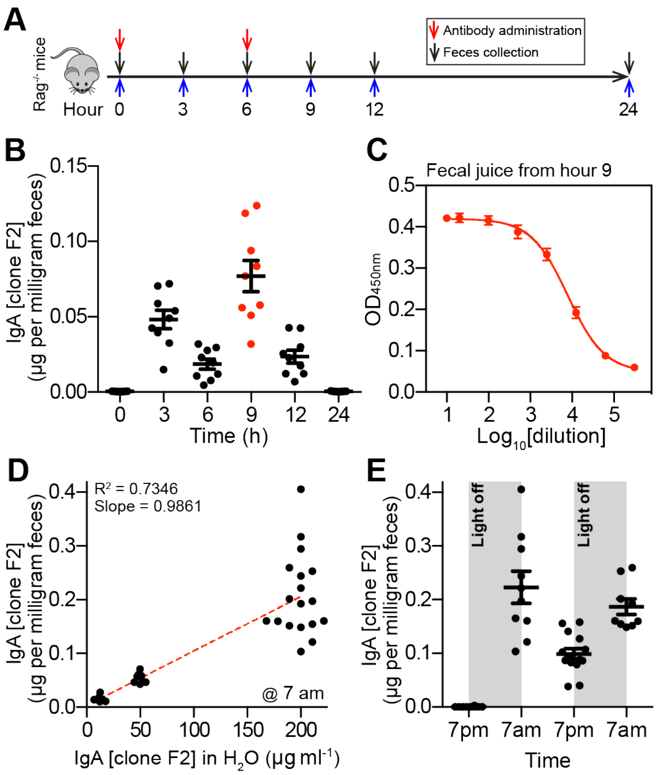


**Fig. S3. Hybridoma-produced monoclonal IgA antibodies still retain binding capacity after passing through intestinal tract, related to fig. 2. (A)** Schematic representation of antibody gavage in Rag$^{-/-}$ mice (100 µg per gavage). **(B)** IgA quantification in the stool of Rag$^{-/-}$ mice at different time points after IgA F2 clone antibody gavage. (**C**) IgA F2 clone binding capacity in the stool of Rag$^{-/-}$ mice at hour 9 after initial gavage. (**D**) Correlation of IgA F2 clone in drinking water (X axis) and stool (Y axis). Mice were given water containing different concentration of IgA F2 clone at 7 pm. The next day morning at 7 am, stool samples were collected and IgA level was quantified by ELISA. (**E**) IgA F2 clone concentration in the stool of mice at different time point. Mice were given water containing 0.2 mg/ml IgA F2 clone at 7 pm. Stool samples were collected every 12 hours in the following 36 hours.

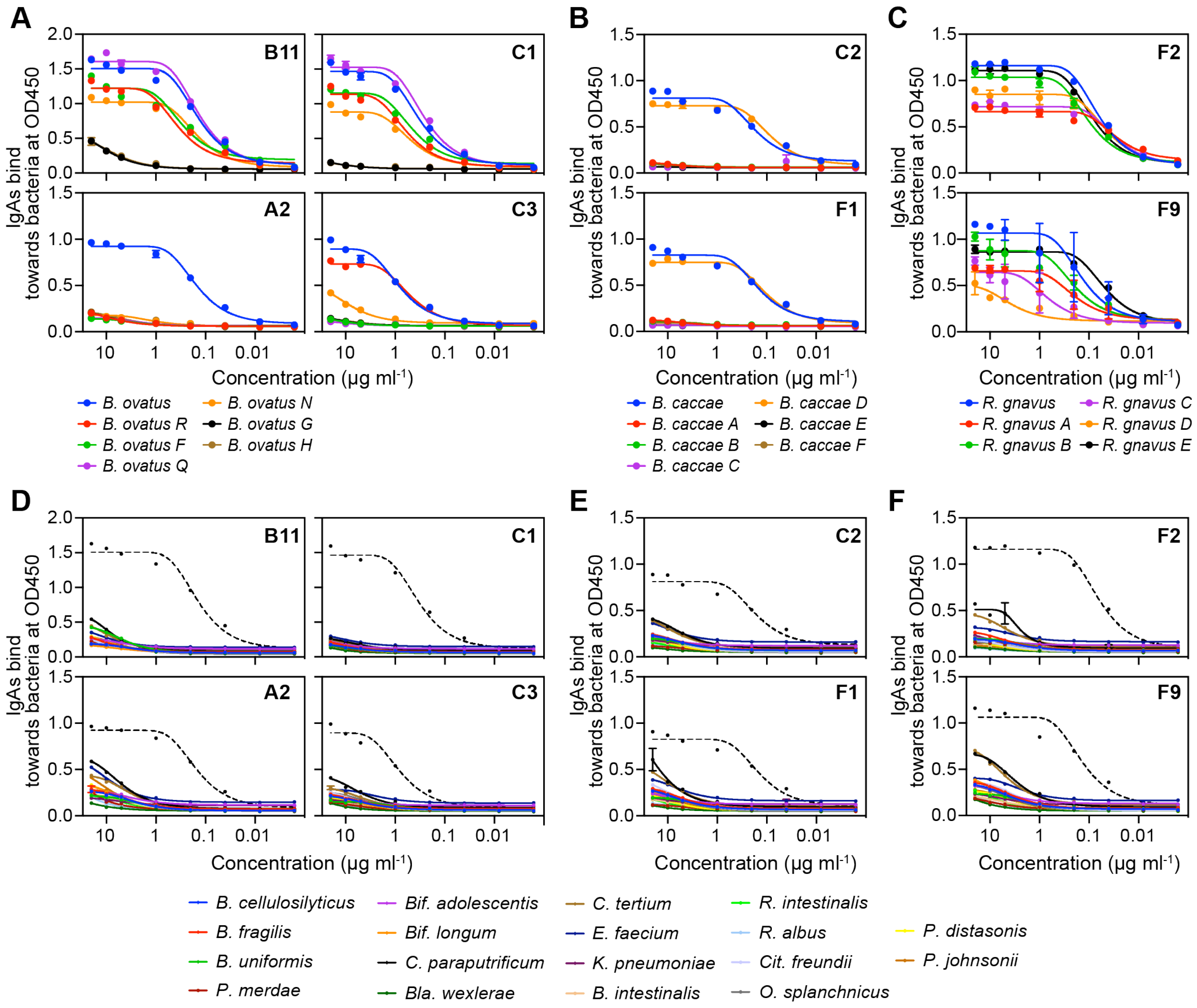


**Fig. S4. Specificity of hybridoma-produced IgA towards different strains and species of bacteria that did not colonized host mice, related to fig. 3.** (**A-C**) ELISA analysis of hybridoma-produced IgA against different strains of *B. ovatus* (**A**), *B. caccae* (**B**) and *R. gnavus* (**C**). Strains of *B. ovatus, B. caccae* and *R. gnavus* obtained from ATCC were used as positive control. (**D-F**) Diverse bacterial species were used to test the cross reactivity of antibodies, which target *B. ovatus* (D), *B. caccae* (E) and *R. gnavus* (F), to gut microbiome by ELISA. $OD_{450}$: optical density at 450 nm. Dotted lines denote positive controls.

**Table S1: Bacterial strains used in this study.**

| Phylum | Species | Strain ID | Cocktail Name | Strain Abbreviation | Sources |
|---|---|---|---|---|---|
| Bacteroidetes | *Bacteroides ovatus* | ATCC8483 | 8M Mix | *B. ovatus* | ATCC |
| Bacteroidetes | *Bacteroides caccae* | ATCC43185 | 8M Mix | *B. caccae* | ATCC |
| Bacteroidetes | *Bacteroides thetaiotaomicron* | ATCCVPI5482 | 8M Mix | *B. theta.* | ATCC |
| Bacteroidetes | *Bacteroides vulgatus* | ATCC8482 | 8M Mix | *B. vulgatus* | ATCC |
| Firmicutes | *Ruminococcus gnavus* | ATCC29149 | 8M Mix | *R. gnavus* | ATCC |
| Firmicutes | *Clostridium bolteae* | ATCC_BAA-613 | 8M Mix | *C. bolteae* | ATCC |
| Actinobacteria | *Collinsella aerofaciens* | ATCC25986 | 8M Mix | *C. aero.* | ATCC |
| Proteobacteria | *Escherichia coli* | ATCC_K-12_MG1655 | 8M Mix | *E. coli* | ATCC |
| Bacteroidetes | *Bacteroides ovatus* | 1001302B_160321_F3 | 8M* Mix | *B. ovatus** | Faith Lab |
| Bacteroidetes | *Bacteroides caccae* | 1001136B_160425_A12 | 8M* Mix | *B. caccae** | Faith Lab |
| Bacteroidetes | *Bacteroides thetaiotaomicron* | 1001136B_160425_G3 | 8M* Mix | *B. theta.** | Faith Lab |
| Bacteroidetes | *Bacteroides vulgatus* | 1001136B_160425_A3 | 8M* Mix | *B. vulgatus** | Faith Lab |
| Firmicutes | *Ruminococcus gnavus* | 1001302B_160321_G6 | 8M* Mix | *R. gnavus** | Faith Lab |
| Firmicutes | *Clostridium bolteae* | 1001302B_160321_D9 | 8M* Mix | *C. bolteae** | Faith Lab |
| Actinobacteria | *Collinsella aerofaciens* | 1001136B_160425_H1 | 8M* Mix | *C. aero.** | Faith Lab |
| Proteobacteria | *Escherichia coli* | 1001136B_160425_B3 | 8M* Mix | *E. coli** | Faith Lab |
| Bacteroidetes | *Bacteroides ovatus* | 1001254J_160919_B1 | N/A | *B. ovatus R* | Faith Lab |
| Bacteroidetes | *Bacteroides ovatus* | BSD3178_07_1175_160815_A10 | N/A | *B. ovatus F* | Faith Lab |
| Bacteroidetes | *Bacteroides ovatus* | BSD2780_06_1687_150420_H2 | N/A | *B. ovatus Q* | Faith Lab |
| Bacteroidetes | *Bacteroides ovatus* | 1001271B_150615_H2 | N/A | *B. ovatus N* | Faith Lab |
| Bacteroidetes | *Bacteroides ovatus* | BSD3448_08_0949_C3 | N/A | *B. ovatus G* | Faith Lab |
| Bacteroidetes | *Bacteroides ovatus* | 1001099B_141217_E5 | N/A | *B. ovatus H* | Faith Lab |
| Bacteroidetes | *Bacteroides caccae* | 1001270I_E9 | N/A | *B. caccae A* | Faith Lab |
| Bacteroidetes | *Bacteroides caccae* | 1001285I_161205_F12 | N/A | *B. caccae B* | Faith Lab |
| Bacteroidetes | *Bacteroides caccae* | BSD3178_07_1176_160815_A7 | N/A | *B. caccae C* | Faith Lab |
| Bacteroidetes | *Bacteroides caccae* | 1001136B_160425_A12 | N/A | *B. caccae D* | Faith Lab |
| Bacteroidetes | *Bacteroides caccae* | 1001283B_150217_161031.161102_F12 | N/A | *B. caccae E* | Faith Lab |
| Bacteroidetes | *Bacteroides caccae* | D43.t1_B9 | N/A | *B. caccae F* | Faith Lab |
| Firmicutes | *Ruminococcus gnavus* | BSD3440_0968_E11 | N/A | *R. gnavus A* | Faith Lab |
| Firmicutes | *Ruminococcus gnavus* | 1001287H_G9 | N/A | *R. gnavus B* | Faith Lab |
| Firmicutes | *Ruminococcus gnavus* | 1001302B_160321_G6 | N/A | *R. gnavus C* | Faith Lab |
| Firmicutes | *Ruminococcus gnavus* | BSD2780_74_G6 | N/A | *R. gnavus D* | Faith Lab |
| Firmicutes | *Ruminococcus gnavus* | D31.t1_F7 | N/A | *R. gnavus E* | Faith Lab |
| Bacteroidetes | *Bacteroides cellulosilyticus* | 1001283B_150304_161114_B8 | N/A | *B. cellulosilyticus* | Faith Lab |
| Bacteroidetes | *Bacteroides fragilis* | J1101437_171009_C3 | N/A | *B. fragilis* | Faith Lab |
| Bacteroidetes | *Bacteroides uniformis* | 1001275B_160808_C2 | N/A | *B. uniformis* | Faith Lab |
| Actinobacteria | *Bifidobacterium adolescentis* | 1001099B_141217_F6 | N/A | *Bif. adolescentis* | Faith Lab |
| Actinobacteria | *Bifidobacterium longum* | BSD2780_12_0874_150323_E3 | N/A | *Bif. longum* | Faith Lab |
| Proteobacteria | *Citrobacter freundii* | 1001302B_160321_C4 | N/A | *Cit. freundii* | Faith Lab |
| Firmicutes | *Clostridium paraputrificum* | 1001275B_160808_B2 | N/A | *C. paraputrificum* | Faith Lab |
| Firmicutes | *Clostridium tertium* | BSD3178_07_1175_160912_G10 | N/A | *C. tertium* | Faith Lab |
| Firmicutes | *Enterococcus faecium* | J1101004_170508_D1 | N/A | *E. faecium* | Faith Lab |
| Proteobacteria | *Klebsiella pneumoniae* | 1001283B_150209_150212_F4 | N/A | *K. pneumoniae* | Faith Lab |
| Bacteroidetes | *Odoribacter splanchnicus* | 1001283B_150304_161114_H6 | N/A | *O. splanchnicus* | Faith Lab |
| Bacteroidetes | *Parabacteroides distasonis* | 1001283B_150217_161031_H2 | N/A | *P. distasonis* | Faith Lab |
| Bacteroidetes | *Parabacteroides johnsonii* | DSMZ_18315 | N/A | *P. johnsonii* | DSMZ |
| Bacteroidetes | *Parabacteroides merdae* | 1001283B_150209_150212_D10 | N/A | *P. merdae* | Faith Lab |
| Firmicutes | *Blautia wexlerae* | 1001283B_150217_161031_B5 | N/A | *B. wexlerae* | Faith Lab |
| Bacteroidetes | *Bacteroides intestinalis* | DSMZ_17393 | N/A | *B. intestinalis* | DSMZ |
| Firmicutes | *Roseburia intestinalis* | J1101437_171009_C5 | N/A | *R. intestinalis* | Faith Lab |
| Firmicutes | *Ruminococcus albus* | 1001217B_150727_F5 | N/A | *R. albus* | Faith Lab |

**Table S2: VDJ sequences of IgA hybridoma clones derived from LP B cells of gnotobiotic mice.**

| Clone | VH | DH | JH | CDR3 sequence | Target Bacteria* |
|---|---|---|---|---|---|
| A2 | V5-16*01 | D4-1*01 | J4*01 | gcaagagctcccttacggggggctatggactac | *B. ovatus* |
| B6 | V1-82*01 | D2-1*01 | J1*03 | gcaagacgctatggtaactactggtacttcgatgtc | All 8 bacterial strains |
| B11 | V1-11*01 | D1-1*01 | J2*01 | ggaagggggaagtttttgactac | *B. ovatus* |
| C1 | V1-11*01 | D3-3*01 | J2*01 | ggaagggggagatctttgactac | *B. ovatus* |
| C2 | V14-3*01 | D2-4*01 | J3*01 | gctattgattacgacgggtttgcttac | *B. caccae* |
| C4 | V5-17*01 | D2-1*01 | J1*03 | gcaaggtatggtaactactggtacttcgatgtc | *B. ovatus and R. gnavus* |
| C8 | V1-62-2*01 or V1-71*01 | D3-3*01 | J3*01 | gcaagacacgaagaaggggacgttttgcttac | N/A** |
| D4 | V5-16*01 | D4-1*01 | J4*01 | gcaagagctcccttacggggggctatggactac | *B. ovatus and R. gnavus* |
| F1 | V14-3*01 | D2-4*01 | J3*01 | gctattgattacgacgggtttgcttac | *B. caccae* |
| F2 | V5-4*01 | D1-1*01 | J2*01 | gcaagagccccttactacggttactactttgactac | *R. gnavus* |
| F3 | V1-66*01 | D3-1*01 | J2*01 | gcaagatgcttggactactttgactac | N/A** |
| F11 | V1-82*01 | D2-1*01 | J1*03 | gcaagacgctaatggtaactactggtacttcgaatgtctg | All 8 bacterial strains |
| G7 | V6-6*01 | D1-1*01 | J3*01 | accaggaactacggtagtaccccgtttgcttac | *B. ovatus and R. gnavus* |

* All bacterial strains were from public repositories (see table S1)

** Do not bind any of the 8 tested bacterial strains or common antigens.
